# Multiplexed microfluidic chip for cell co-culture

**DOI:** 10.1101/2022.03.25.485382

**Authors:** Craig Watson, Chao Liu, Ali Ansari, Helen C. Miranda, Rodrigo A Somoza, Samuel E. Senyo

## Abstract

Paracrine signaling is challenging to study in vitro, as conventional culture tools dilute soluble factors and offer little to no spatiotemporal control over signaling. Microfluidic chips offer potential to address both of these issues. However, few solutions offer both control over onset and duration of cell-cell communication, and high throughput. We have developed a microfluidic chip designed to culture cells in adjacent chambers, separated by valves to selectively allow or prevent exchange of paracrine signals. The chip features 16 fluidic inputs and 128 individually-addressable chambers arranged in 32 sets of 4 chambers. Media can be continuously perfused or delivered by diffusion, which we model under different culture conditions to ensure normal cell viability. Immunocytochemistry assays can be performed in the chip, which we modeled and fine-tuned to reduce total assay time to 1h. Finally, we validate the use of the chip for co-culture studies by showing that HEK293Ta cells respond to signals secreted by RAW 264.7 immune cells in adjacent chambers, only when the valve between the chambers is opened.

## Introduction

Paracrine signaling is central to countless biological processes, from development to wound healing and tissue homeostasis. It is challenging to study, however, as conventional in vitro tools are limited in their ability to recapitulate paracrine signaling in a physiological relevant manner, diluting signals and providing little control over spatial and temporal aspects of signaling. Several microfluidic chips have recently emerged to address these issues with various designs (1, 2). A key advantage of microfluidic versus traditional cell culture tools is limiting dilution of cell-secreted signals. Multiple studies have demonstrated that the volume of media in which cells are cultured can affect differentiation, epithelialto-mesenchymal transition and other aspects of cell behavior (3–5). While the specific impact of low or high volume varies between cell types, microfluidic culture systems offer a straightforward way to control concentration of autocrine and paracrine signals by adjusting media supply rate (6–8). Microfluidic chips to study cell-cell signaling can be broadly sorted into two categories: “always-on” devices, where the cells are in contact with each other from the moment of seeding until the end of the experiment, and “on-off” or “selective” devices, which offer a way to enable or disable the exchange of paracrine signals between cells at any given time. Selective co-culture can be achieved by tuning surface wettability (9) or including valves (10–13), with the latter strategy offering the most flexibility over initiation and duration of coculture as well as allowing multiple conditions to be studied in parallel. Few solutions, however, offer both control over initiation of co-culture and high throughput.

Here, we present a microfluidic chip for mammalian cell culture featuring valves to enable selective co-culture as well as multiple simultaneous experimental conditions with 128 individually-addressable chambers. The ability to deliver media by perfusion or diffusion makes the chip suitable for a large range of co-culture studies, and the use of a novel plastic coverslip-thickness substrate allows high resolution imaging and efficient cell attachment. We validate the applicability of our chip to co-culture studies by demonstrating response of HEK293Ta cells to cytokines secreted by RAW 264.7 monocyte/macrophage-like cells in neighboring chambers. We further characterize the chip by modeling nutrient consumption under different media supply conditions, and detail immunocytochemistry protocols optimized for use in the chip.

## Results and Discussion

### Chip design

The chip (Fig. 1) is made by standard two-layer soft lithography with push-down valves (14, 15). The chip features 16 fluid inputs, 32 identical cell culture units and 2 outputs. Each unit is composed of a rinse channel and 4 parallel chambers. Neighboring chambers are connected by short channels which allow for diffusion between them and which can be closed with a valve. Cells can be cultured in all 4 chambers, or in the central 2 chambers with the outer 2 chambers used as media reservoirs. In the latter case, the central valve (between chambers B and C, Fig. 1B) can be opened or closed to selectively allow or prevent exchange of paracrine signals between the two adjacent cell populations. Two control lines are used for this central valve, each connected to 16 units of the chip, therefore making it possible to conduct co-culture and isolated culture simultaneously in different units of the chip.

**Fig. 1.**
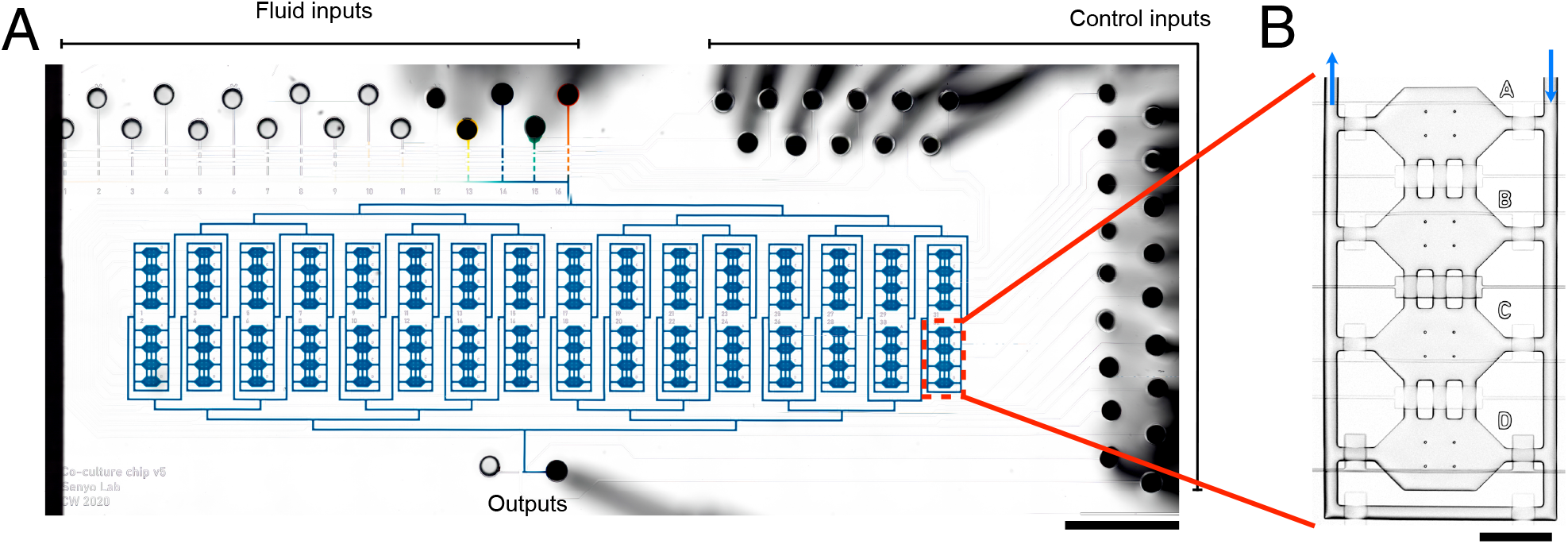
**A)** Stitched brightfield image of chip with food dye in all channels and chambers. 16 inputs combine into one channel which then splits off into 32 branches of equal length (and thus fluidic resistance) to supply the 32 units. The channels downstream of the units recombine and split off into two outputs. The chip measures approximately 60 mm x 20 mm, fitting on a standard microscope slide. Scale bar: 5 mm **B)** Detail of one unit, comprising 4 chambers and a rinse channel. Push-down valves are placed at the inlet and outlet of each chamber and channel, and between the chambers. The valves are common to all units, except for the valve between chambers B and C which is connected to one of two different control lines, to allow the valves to be closed on half of the chip and open on the other half. Arrows indicate flow direction. Scale bar: 500 μm

The 16 inputs are addressed by an 8-valve flat multiplexer, and combine into one channel which then splits to the 32 units with a branched, 10-valve multiplexer (16). Each branch is of equal length, yielding equal flow rate to each unit regardless of the number of units addressed at a time. The multiplexing results in each of the 128 chambers being individually addressable while requiring only 29 control lines for the entire chip (Fig. S1).

A custom graphical user interface was designed for the chip, with pre-programmed multiplexer settings, input labels and pressure control (Fig. S2A)(17), thus abstracting complexities of chip operation and allowing intuitive control. Though the software is specifically designed for our pneumatic controller, it is open source and can easily be modified to communicate with other hardware setups.

### Substrate

The PDMS chips were sealed in three different ways: to glass slides (after plasma treatment), to PDMS-coated glass slides, or to coverslip-thickness plastic slides. The plastic slides improve upon common glass/PDMS substrates by allowing high-resolution imaging without the brittleness of glass coverslips, as well as enabling long-term live culture and imaging with no artifacts. When using a glass slide as substrate, water vapor diffusing from the channels through the PDMS gets trapped on the surface of the slide, forming droplets around channels, including below them (in the case of a PDMS-on-glass substrate). Plastic coverslip-thickness substrates mitigate this issue thanks to their vapor permeability (Fig. S3).

Furthermore, cells attachment to plastic slides is as efficient as on conventional polystyrene culture substrates, and better than on PDMS (Fig. S4). A wide variety of cells have been cultured in the chip, from cell lines to primary cells, with support for various optical read-outs including high-magnification immunocytochemistry images (Fig. 2). Long term experiments are also possible, as we have cultured cells for up to 14 days. While the chip was used with all three substrates, plastic slides were chosen for the majority of experiments.

**Fig. 2.**
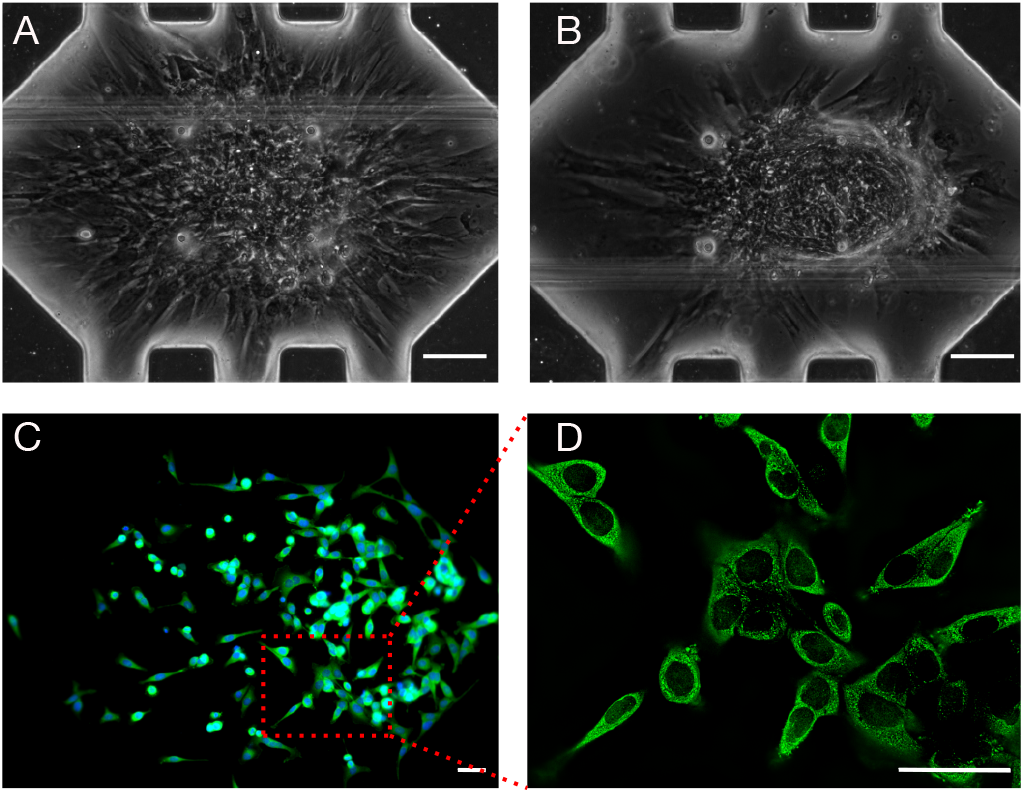
The co-culture chip supports a variety of different cells and experimental read-outs. **A**,**B)** Primary human mesenchymal stem cells (MSCs) after 7 days of culture in low-glucose DMEM (A) or TGF-β-supplemented DMEM (B). Depending on media composition, MSCs will either spread out or form micro-masses. **C)** HeLa cells after 2 days of culture, fixed and stained for p65. **D)** close-up of the same chamber, taken with 40x oil immersion objective and deconvolved. Scale bars: A,B: 100 μm; C,D: 50 μm

One limitation of plasma-bonded plastic and glass substrates is a restriction in channel design: since plasma treatment stiffens PDMS (18, 19), higher pressures are necessary to close valves completely; these high pressures increase likelihood of delamination between the flow and control layer and of leaks between the inlet holes and needles. In all cases, channels must be tall enough to prevent collapse, but short enough to enable reliable valve operation. The same chip, when assembled without plasma treatment on a PDMS-coated substrate, could support channel heights of up to 30 μm versus 22 μm when plasma-bonded. Taller channels (and chambers) are desirable to avoid restricting large cells, thus a trade-off between channel height and substrate choice must be considered depending on the application.

### Media supply

The chamber layout allows several media delivery approaches: perfusion through the culture chambers, or diffusion from the adjacent chamber which serves as a reservoir. In both cases, the flow can be continuous or “pulsed”, where flow is started and stopped periodically to prevent excessive dilution of cell-secreted signals. We mainly used a pulsed diffusion scheme (10) where media are delivered to an adjacent chamber, flow is stopped, and the valve between the two chambers is lifted to allow diffusion of metabolites and waste products. The valve is then closed, fresh media are flowed to the reservoir and the cycle is repeated (Fig. 3A).

**Fig. 3.**
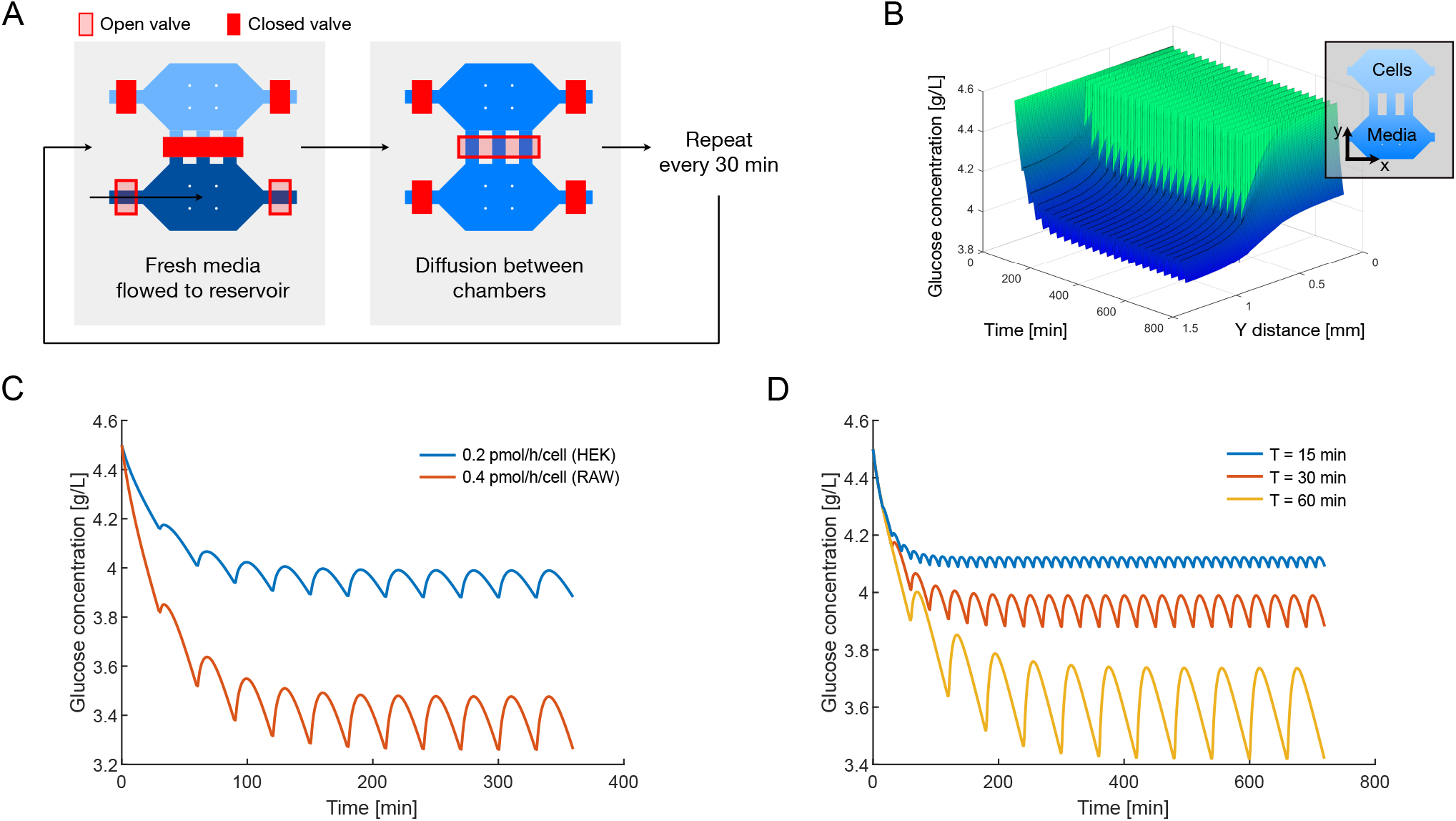
Glucose diffusion and consumption model. **A)** Schematic of media supply by diffusion. Medium is flowed into the reservoir adjacent to the culture chamber. The valve between the chambers is then opened to allow diffusion with no shear stress on the cells. This is repeated approximately every 30 minutes, depending on cell types. **B)** Glucose concentration in chamber vs time. Concentration over x axis rapidly becomes homogeneous due to the small distance between channels, therefore results are averaged over the x axis for better visualization. Media changes are modeled by resetting the concentration in the reservoir and channels to 4.5 g/L. **C)** Impact of glucose consumption rate on the concentration at end of the chamber farthest from the media reservoir. Media supply period is set to 30 min. **D)** Impact of media supply period T on glucose concentration at far end of chamber, with consumption rate set to 0.2 pmol/h/cell.

While this scheme minimizes dilution of cell-secreted factors, the pulse frequency must be chosen to allow sufficient supply of fresh media. To ensure that our chamber design would allow for sufficiently rapid exchange, we modeled the diffusion and consumption of glucose with pulsed supply. The model accounts for chamber geometry, cell density, glucose consumption per cell and media supply frequency, and outputs glucose concentration over time and position within the chambers (Fig. 3B).

Since glucose consumption varies between cell types, we compared the profile of HEK293 and RAW264.7 cells based on reported consumption rates, with values of 0.2 and 0.4 pmol/h/cell (Fig. 3C) (20, 21). Next, in order to lower the chance of glucose levels negatively impacting cell function, we compared media resupply periods to keep concentration within ~70% of its maximum, and chose 30 to 60 minute periods for the cells presented here (Fig. 3D).

The model provides only a limited view of cell metabolism and does not include factors such as media pH or oxygenation, but provided sufficient information to guide the design of chamber geometries and initial values for media supply period, which we then tested empirically. In actual experiments, supply periods of 30 to 60 minutes were used for all cell types tested. The model can further be easily altered to visualize secretion and dilution of cell-secreted signals, by setting the appropriate diffusion coefficient and secretion rate of a signal of interest (provided they are known or can be estimated). For more in-depth studies modeling both secretion and uptake of a signal with pulsed media supply, the geometry can be extended to include all 4 chambers.

### Immunocytochemistry

Immunocytochemistry was performed for various assays in the chip. Although the overall protocol is similar to that in conventional multiwell plates, the limited chamber volumes result in rapid depletion of antibodies, and thus solutions must be perfused through the chambers rather than simply incubated. On the other hand, perfusion offers the opportunity to greatly speed up the assay. In order to optimize the protocol to reduce assay time and reagent usage while increasing signal intensity, we built a computational model of the convection, diffusion and binding of antibodies to cells in the chamber. The chamber was modeled in 2D (length × height), and the problem was simplified to consider antibody binding as a surface reaction on the bottom of the chamber, neglecting the height of cells and diffusion through cytoplasm / nucleus. The full COMSOL model including all parameters is included in supplementary file 3.

We first evaluated the impact of bulk antibody concentration on binding rate, using mean surface concentration of antibodies over time as the output of interest (Fig. 4A). We then used time to 90% of saturation as a summary to evaluate the effect of different parameters on assay time (Fig. 4B). Next, we looked at the effect of flow rate on assay time (Fig. 4C), which clearly indicates the benefit of flow versus static incubation, with diminishing returns once the flow rate permits quasi-instant replenishment of bulk antibodies close to the reaction surface.

**Fig. 4.**
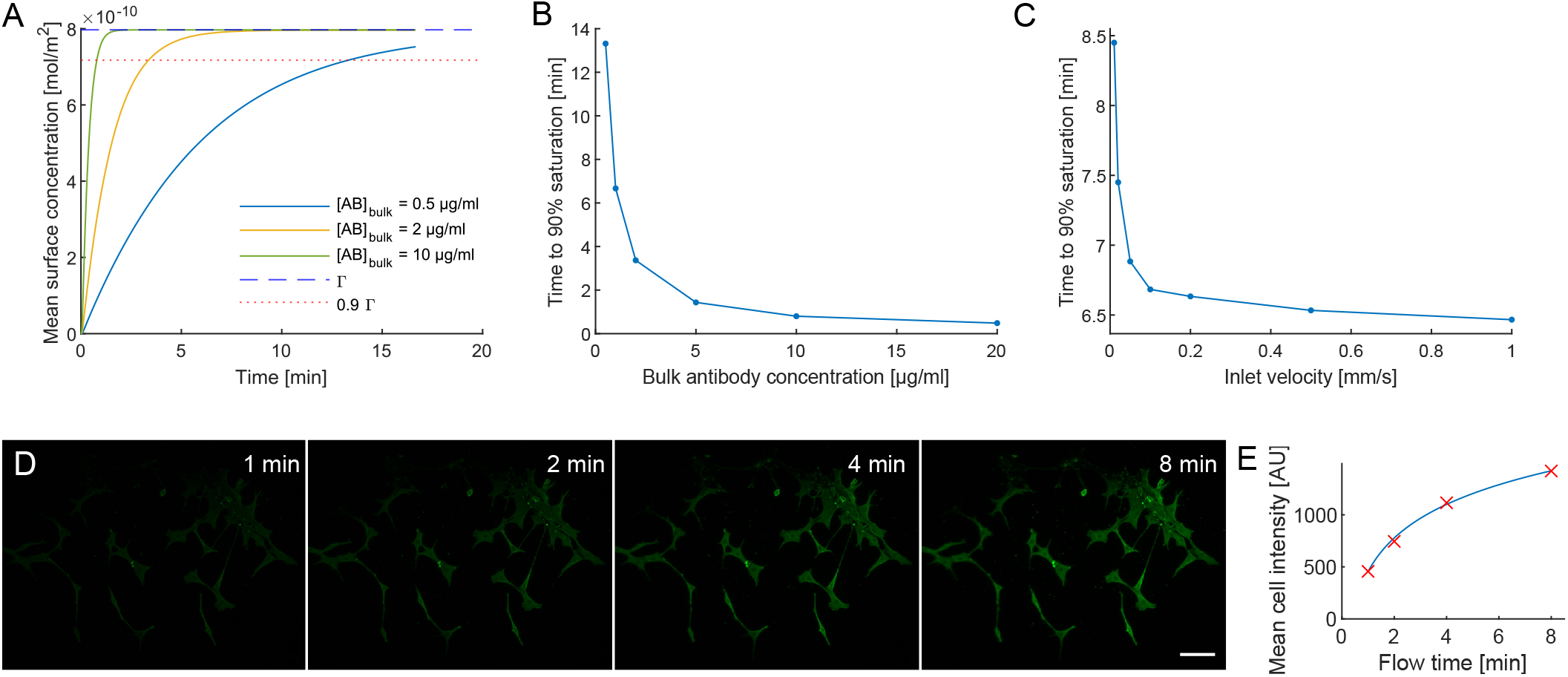
Immunocytochemistry model. **A)** Impact of bulk antibody concentration on concentration of surface-bound antibodies. Dashed and dotted lines represent surface concentration of antibody binding sites (i.e. saturation) and 90% saturation respectively. Inlet velocity was set to 0.1 mm/s. **B)** Summary of the relationship between bulk antibody concentration and the time to reach 90% of saturation. **C)** Impact of antibody solution flow rate on binding time, with bulk concentration set to 1 μg/mL. Large gains are made by overcoming the diffusion limitation, with diminishing returns as flow rate is increased. **D)** Experimental validation: secondary antibody (at 10 μg/mL) flowed into chamber with 3T3 cells after fixation and flow of anti-vimentin primary antibodies. Scale bar: 100 μm. **E)** Mean intensity of the cells (excluding background areas) in D, revealing brightness increase over time consistent with model.

The model was built as a convection-diffusion-reaction simulation with the reaction occurring at the bottom surface of the chamber. This does not correspond exactly to the real case where cells have a non-zero height and antibodies may bind either on the surface or interior of cells. Obtaining diffusion coefficients through permeabilized cells as well as estimating antibody binding capacity would require extensive experimentation for little added gain; therefore, the model was simplified as described and used to estimate the effect of different parameters on the outcome of staining, without fully recapitulating the process. We then conducted follow-up experiments to fine-tune actual concentrations and flow times of different reagents in the chip.

Based on the results of this model, we increased antibody concentration 4-fold compared to recommended protocols for multiwell plates, and perfused all reagents at 4-6 kPa. Perfusion time for secondary antibodies was adjusted based on direct visualization with time lapse imaging. While validation in the chip did not permit quantitative comparison between conditions such as antibody concentration or flow rates, the profile in any given condition matched the computational model well (Fig. 4D,E). Overall time for the complete fixation and staining protocol for 32 chambers (1 chamber in each of the 32 units) was reduced to approximately 60 minutes (Sup. Note 1), with the entire protocol carried out automatically after initial connection and degassing of the inputs. Less than 50 μL of each reagent was sufficient.

### Co-culture

To validate the use of the chip for coculture studies, we made use of the TNFα-mediated NF-κB activation model. When activated with LPS and/or IFNγ, macrophages secrete inflammatory cytokines including TNFα that can activate the NF-κB pathway in other cells (Fig. 6A) (22–24). We generated a stable HEK293 cell line expressing a fluorescent (mKate2) reporter for NF-κB (25); after verifying robust reporter activation in the dish when exposed to TNFα (Fig. S5), we conducted a simple study in the chip to make use of this pathway by selectively exposing HEK293 reporter cells to inflammatory cytokines secreted by RAW 264.7 monocyte/macrophage-like cells.

A total of 8 experimental groups with 4 replicates each were established (Fig. 5). HEK293 and RAW 264.7 were seeded in adjacent chambers of 16 units of the chip; in 8 of these units, RAW cells were treated with LPS while the RAW cells in the 8 other units were kept untreated. In the remaining 16 units, HEK293 were cultured alone either in normal medium or in LPS-or TNFα-supplemented media. After culturing cells overnight, the valve between HEK and RAW chambers was lifted in half of the units to allow exchange of paracrine signals.

**Fig. 5.**
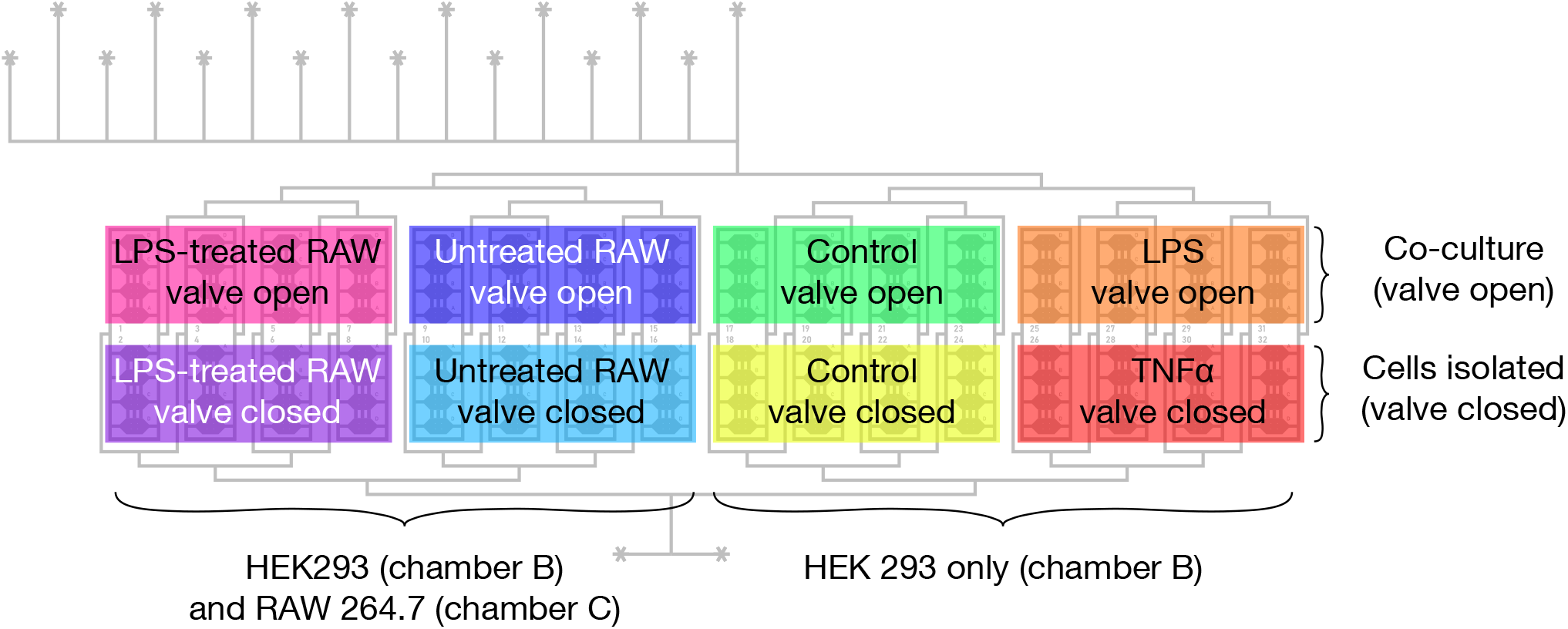
Schematic of the experimental groups in the NF-κB demonstration experiments. HEK293 were seeded in chamber B of all units; RAW were seeded in chamber C of units 1-16. The next day, RAW cells in 1-8 were briefly treated with LPS. Then, the valve between chambers B and C in odd-numbered units (top half of chip) was opened to allow exchange of paracrine signals. Normal medium was delivered to all units, except four which received LPS-supplemented medium (top right); and four which received TNFα-supplemented medium (bottom right). This yielded 8 experimental groups with 4 technical replicates each.

24h later, we observed strong activation of the NF-κB reporter in the HEK cells when exposed to RAW cells or to TNFα, but not in any other conditions (Fig. 6B-D). No activation was observed in HEK cells cultured adjacent to RAW cells with the valve closed, confirming that media were not able to diffuse between those chambers. There was no difference between valve open and closed conditions for HEK293Ta cultured alone, suggesting that the slight difference in media volume had no effect on reporter expression. The fraction of positive cells in co-culture (valve open) conditions matched the TNFα positive control. No difference between the effects of LPS-treated and untreated RAW cells was observed, indicating that inflammatory cytokines were secreted in both conditions – while this response even with non-treated macrophages was unexpected, we did not seek to further modulate macrophage activation (for example by treatment with IL-10 (22)) for the purpose of this validation study.

**Fig. 6.**
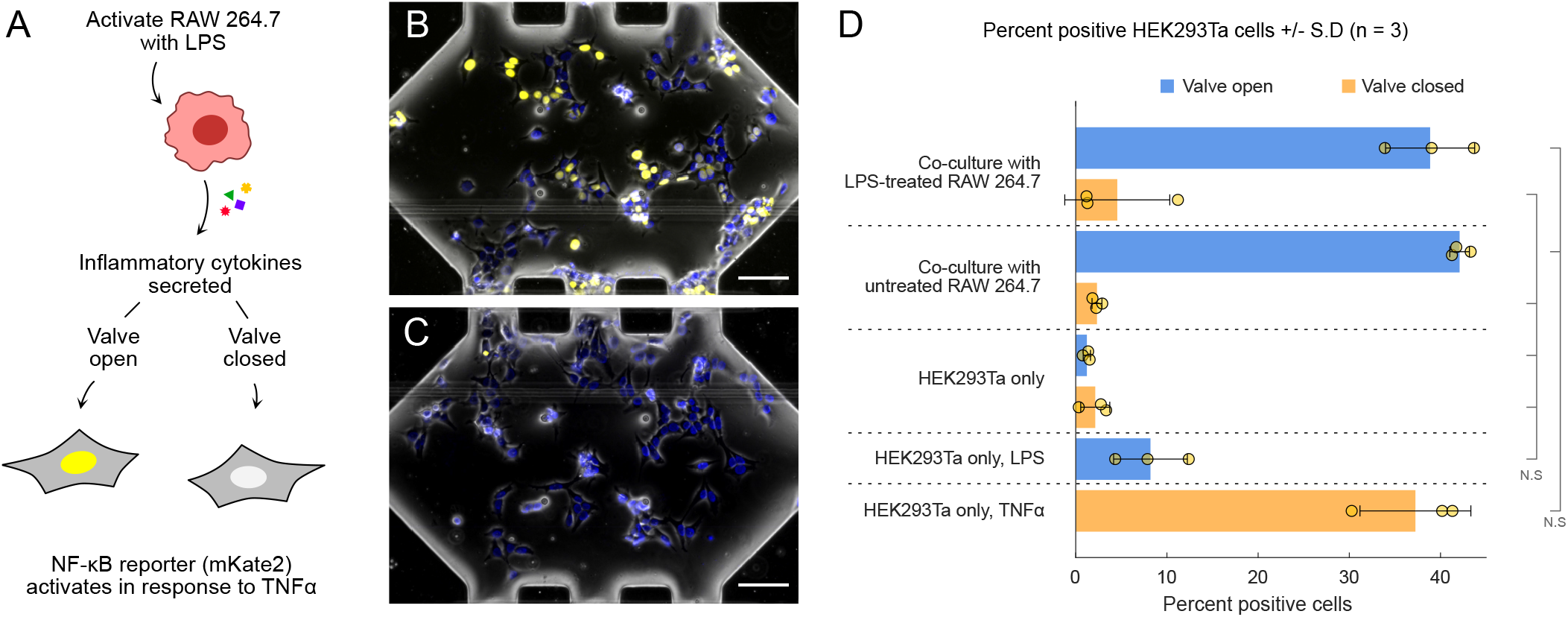
Co-culture validation. **A)** RAW 264.7 and HEK293 (transduced with a fluorescent NF-κB reporter) are cultured separately in neighboring chambers. LPS is delivered to the RAW cells, activating them to stimulate production of inflammatory cytokines such as TNFα. The valve between the RAW and HEK chambers is then opened in half of the units of the chip, allowing cytokines to diffuse towards the HEK cells. **B, C)** Representative images of HEK293 chambers in the valve open (B) and valve closed (C) conditions. Yellow: mKate2; blue: Hoechst; gray: phase contrast. Scale bars: 100 μm. **D)** Quantification, showing mean number of positive cells in each condition. 1 dot = 1 experiment; 4 units (technical replicates) per group. HEK293Ta were either co-cultured with RAW 264.7 (top 4 groups) or alone (bottom 4 groups). Isolated HEK293Ta were supplied with either normal media or media supplemented with LPS (100ng/mL) or TNFα (100 ng/mL). Groups are significantly different (p < 0.01) except where noted (N.S: not significant). All comparisons shown in Fig. S6.

These experiments demonstrate that co-culture and isolation can be studied simultaneously on the chip, with cells in otherwise identical conditions; that co-culture can be initiated at arbitrary timepoints; and that experiments with multiple experimental groups can be carried out with ease. Here, media supply (including periodic switching between control and supplemented media, with rinse steps) was performed autonomously.

## Materials and Methods

### Mold fabrication

100 mm silicon wafers (UniversityWafer, test-grade silicon) were used to create negative molds for the control and flow layers. Flow layer wafers were sputtercoated with 5 nm of titanium, then AZ9260 photoresist (MicroChemicals) was spin-coated to a thickness of 22 μm and patterned with a transparency photomask (CAD/Art services, Brandon OR USA). Following development, the wafers were heated to 125°C for 2 min on a hot plate to obtain the rounded profile necessary for valve operation (10). Control layer wafers were spin-coated with SU-8 2025 (Kayaku Advanced Materials, Westborough MA USA) to a thickness of ~30 μm, and patterned with a transparency photomask. Photomasks were designed with Autodesk AutoCAD; full designs are included in supplementary file 1. Following photolithography, both control and flow layer wafers were coated with 2 μm Parylene C to promote reliable PDMS demolding, with the same coater and process as described (26). Molds were used up to 73 times before failure.

### Chip fabrication

PDMS chips were cast using 2-layer soft lithography techniques (14). Sylgard 184 base and curing agent (Dow 1317318) were mixed at a 5:1 base:crosslinker ratio, poured onto the control wafer and partially cured on a hot plate, covered, for 30 min at 80°C. Control inlets were then punched using a 1 mm biopsy punch (Electron Microscopy Sciences 69039-10) mounted on a drill press stand. 20:1 PDMS was spin-coated onto the flow layer at 850 rpm for 60 s, left to reflow for 15 min, then partially cured on a hot plate for 6 min at 80°C. The control layer chips were then aligned onto the flow layer manually under a stereo microscope and the assembly baked for a further 30 min. Chips were then removed from the wafer, and inlet holes punched. Plastic slides (Ibidi 10813) were functionalized with O_2_ plasma (30 W / 30 s) followed by incubation in 1% v/v (3-Aminopropyl)triethoxysilane (APTES) (Sigma-Aldrich 440140) in water for 20 min, rinsed then blow dried with nitrogen (27). The PDMS chips were then plasma treated (22 W / 30 s), bonded onto the treated plastic slides, and baked for 1 h at 80°C. Chips were kept at room temperature for a minimum of two weeks before use to prevent excessive permanent collapse of valves. For glass-substrate chips, both PDMS and glass slide were plasma treated (22 W / 30 s) simultaneously and bonded, before baking for 1 h at 80°C. Alternatively, chips were bonded to a slide by spin-coating 20:1 PDMS on a glass slide (2300 rpm / 30 s), partially curing by baking at 80 °C for 12 min, then assembling the chips onto the slide and baking for one hour more.

### Metabolic model

Glucose metabolism was modeled using MATLAB. Live scripts including an in-depth explanation of the model are provided in supplementary file 2. The chamber geometry was modeled in 2D, with two different domains corresponding to the media reservoir and the culture chamber. A simple diffusion-consumption model was used, with the change in glucose concentration over time defined as:

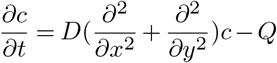

Where *D* is the diffusion coefficient of glucose, and *Q* is the consumption rate. *Q* = 0 in the media reservoir and channels (diffusion only, no consumption) and *Q* = *Q*_0_ in the cell chamber. Consumption was modeled as constant, i.e. not depending on the available glucose concentration. To account for periodic media supply, glucose concentration in the reservoir was reset to 4.5 g/L at specified intervals. Consumption rate in the chamber was calculated from a given rate per cell, and a given cell density per unit area.

### Immunocytochemistry model

A convection-diffusionreaction model was built with COMSOL Multiphysics 5.5 (supplementary file 3). The problem was simplified by considering the antibody binding region as the flat, bottom surface of the channel (neglecting the height of cells, and diffusion through the cytoplasm) (28–32). Approximations were made for the density of antigens on the surface based on antibody binding capacity reported for cell surface antigens in the context of flow cytometry (33–35). We chose a value of 1×10^5^ sites/cell. Antibody association and dissociation rates (*k*_*on*_ and *k*_*off*_) were approximated as 1×10^5^ s^−1^M^−1^ and 1×10^−5^ s^−1^ respectively (36). The surface reaction was defined as:

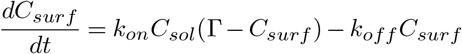

Where *C*_*surf*_ and *C*_*sol*_ are the concentrations of antibodies on the surface and in solution, respectively, and Γ is the total surface concentration of binding sites, calculated from the antibody binding capacity and the number of cells on the surface. The full list of parameters and variables is documented in the model. Both 2D and 3D models were constructed and, having confirmed that the results were similar, the 2D model was used for all subsequent studies for faster performance. Output of the COMSOL model was processed in MATLAB to produce plots.

#### Validation

Fixation, permeabilization, blocking and primary antibody steps were performed identically for all conditions. Time lapse images were acquired during perfusion of secondary antibody solution. Fluorescence intensity was measured using CellProfiler. The final image was first used to define a mask encompassing all cells; intensity was then measured within these regions for each image corresponding to the 1, 2, 4 and 8 minute timepoints.

### Chip operation

Chips were operated using our previously described automated microfluidics control system (17), updated with a graphical user interface (GUI) customized to this chip design (Fig. S2A). Chips were imaged using an Olympus IX81 microscope equipped with a motorized stage (Prior Scientific, Cambridge, UK), and heated enclosure and stagetop CO_2_ incubator (In Vivo Scientific, Salem SC USA), in which the chips were kept at 37°C / 5% CO_2_ for the duration of the experiments (Fig. S2B). Tygon tubing (McMaster-Carr 5155T11) was used for the control lines, and PTFE tubing (Cole-Parmer EW-06417-31 or EW-06417-21) was used for reagents, inserted into the chip inlets directly or using stainless steel needles manually extracted from 20 gauge luer lock syringe needles (McMaster-Carr 75165A677). For long duration experiments, culture media were delivered through a bottle fitted with a delivery cap (Cole-Parmer EW-12018-54) and a bubble trap (PreciGenome PG-BT-STD25UL). All tubing and needles were autoclaved before use. Following pressurization of the control layer to ~150 kPa, the flow channels were filled with sterile PBS and briefly washed with 70% ethanol, then rinsed with PBS.

### Cell culture

HeLa cells were kindly provided by Horst von Recum, and RAW 264.7 by Andrei Maiseyeu; HEK293Ta were obtained from GeneCopoeia. All cells described here were cultured in DMEM supplemented with 10% FBS and 1% penicillin/streptomycin (Gibco 11995, 10082147, and 15140122).

For HEK293 and HeLa cells, chambers were coated with fibronectin (37). First, Pluronic F-127 (Sigma P2443) at 0.2% w/v in PBS was flowed through all rinse channels for 30 min and rinsed with PBS for 5 min, to prevent cell adhesion outside of the chambers. Then, fibronectin (Sigma FC010) at 30 μg/mL was flowed through the culture chambers for 5-15 min and immediately rinsed with PBS for 3 min. Fibronectin coating was not necessary for RAW 264.7, 3T3 and other cells.

Chips were kept in CO_2_ for a minimum of 2 h prior to cell loading, to allow CO_2_ levels to equilibriate within the PDMS and prevent cell death due to excess basicity.

Cells were resuspended to a density of 5-10 million cells/mL in DMEM supplemented with 16-18% v/v OptiPrep (Sigma D1556) to maintain cells at neutral buoyancy and thus prevent sedimentation in the inlet (which would lead to uneven cell counts in the chambers)(38–40). 5 μL of the cell suspensions were aspirated into 24G PTFE tubing using a pipette inserted into or over the other end of the tubing. The tubing was then connected to the chip, and the suspensions were flowed into multiple chambers at a time (either all 32 or in smaller groups of 4-8 chambers).

Media were supplied to the cells primarily by pulsed diffusion, as described in Fig. 3. Media supply frequency was determined based on cell type, either using our glucose consumption model or empirically. Alternatively, media were perfused directly through the chambers at low flow rates, completely replacing media in the chambers every 60 min. In the case of perfusion, cells were first allowed to attach for 30-60 mins before starting media supply routines.

### Lentiviral transduction

The reporter plasmid was a gift from Timothy Lu (Addgene plasmid #82024) (25). Plasmid-containing bacteria were selected with carbenicillin (Fisher Scientific BP26481) at 100 μg/mL. Plasmid DNA was then isolated with ZymoPURE II plasmid midiprep kit (Zymo Research D4200) and stored at −80°C until use. HEK-293Ta cells were used to produce lentivirus as described previously (41). HEK-293Ta and HeLa were then plated at 1×10^4^ cells/cm^2^ and transduced as described previously (41, 42), with a multiplicity of infection of 1. Transduced cells were selected with puromycin (InvivoGen ant-pr-1) for 72 h at concentrations of 2 and 0.5 μg/mL for HEK-293Ta and HeLa respectively. Reporter activation of selected cells was tested in the dish with exposure to TNFα (Abcam ab9642) and LPS (lipopolysaccharides from E. coli; Sigma L8274); HEK-293Ta showed the strongest response to TNFα and were thus selected as responder cells in the co-culture assay (Fig. S5).

### NF-κB reporter assay

Reporter activation was tested similarly in 48-well plates and the microfluidic chip. In multiwell plates, cells were plated and supplemented media were added to the wells the next day. In the chip, HEK and RAW cells were seeded at 6 and 8 million cells/mL respectively as described above, and media delivered by perfusion every 60 mins overnight. LPS-supplemented DMEM (100 ng/mL) was then perfused to RAW chambers for 2 min and incubated for 30 min, then rinsed. Normal and supplemented media were then delivered by pulsed diffusion (60 min intervals) to different units as in Fig. 5.

In both the dish and chip, 24 h after stimulation onset, cells were counterstained with Hoechst 33342 (Abcam ab228551) and imaged. Image analysis was performed using CellProfiler 4 (43) and MATLAB R2021b; the pipeline and code are available in supplementary file 4. Briefly, individual cells were identified from the Hoechst channel, and mKate2 intensity was measured in each cell. For each image, a threshold was defined as the mean + 2 *×* (standard deviation) of the background intensity, and cells above the threshold considered positive. Groups were compared by one-way ANOVA with Tukey post-hoc test, with significance threshold of p < 0.01.

In preliminary studies, time lapse imaging was used to gauge reporter activation timescale, but was not used in experiments with endpoint fluorescence quantification due to excessive photobleaching.

## Conclusions

The co-culture chip provides a versatile platform to perform paracrine signaling studies, as well as other cell culture studies with high degree of multiplexing. Cell populations can be seeded separately and treated with exogenous factors independently before onset of paracrine signaling; many experimental conditions can be evaluated simultaneously, with 32 to 64 independent co-culture units depending on media supply modality, or up to 128 in monoculture.

The chip can support a wide variety of cells and can be used for long-term studies, as we have conducted experiments for up to 2 weeks in the chip. It is particularly well suited to optical read-outs including fluorescent reporters, chemical stains and immunostaining, with high quality and high resolution imaging thanks to the coverslip-thickness substrate. In addition, chip operation is relatively simple given its complexity thanks to the GUI which abstracts the details of the multiplexers and provides a live view of the state of the chip.

Thanks to automation and precise control over reagent delivery timing, microfluidics are especially suited to studying the dynamics of pathways such as NF-κB, as has been done in single populations of cells (37, 44–47); our chip is amenable to extending these types of studies to incorporate cross-talk between different cells. Treatments can also be delivered to one cell population at different intervals before enabling exchange of paracrine signals, to study the time course of pathways that respond to exogenous signals.

## ACKNOWLEDGEMENTS

We thank Laurie Dudik, Ina Martin and Nichole Hoven for assistance with microfabrication, Hari Baskaran for feedback on the models and manuscript, and Horst von Recum and Robert Kirsch for logistical support. This work was supported by the following grants: NIH R25-HL145817 (SES), ADVANCE Opportunity Grant (SES), 5T32HL134622 (AA), R01 NS121374 (HCM), 1P41EB021911 (RS; Case Center for Multimodal Evaluation of Engineered Cartilage), NIH 1 C06 RR12463-01 (CWRU).

## AUTHOR CONTRIBUTIONS

CW developed the microfluidic chip and fabricated molds, built the computational models, performed data analysis, and drafted the manuscript. CW and CL performed the co-culture experiments. CW, CL and AA developed the PDMS-plastic bonding method, fabricated and tested chips, performed preliminary studies and developed cell culture protocols. CL and RAS generated the reporter cell lines. HCM contributed materials and supervised preliminary work. CW and SES designed the co-culture chip and the study. SES supervised the study. All authors reviewed and edited the manuscript.

## References

1. Mario Rothbauer, Helene Zirath, and Peter Ertl. Recent advances in microfluidic technologies for cell-to-cell interaction studies. Lab on a Chip, 18(2):249–270, 2017. ISSN 1473-0189. doi: 10.1039/C7LC00815E.

2. Timothy Quang Vu, Ricardo Miguel Bessa de Castro, and Lidong Qin. Bridging the gap: Microfluidic devices for short and long distance cell–cell communication. Lab on a Chip, 17 (6):1009–1023, 2017. ISSN 1473-0197, 1473-0189. doi: 10.1039/C6LC01367H.

3. Pouria Fattahi, Amranul Haque, Kyung Jin Son, Joshua Guild, and Alexander Revzin. Microfluidic devices, accumulation of endogenous signals and stem cell fate selection. Differentiation, 112:39–46, March 2020. ISSN 0301-4681. doi: 10.1016/j.diff.2019.10.005.

4. Amranul Haque, Pantea Gheibi, Yandong Gao, Elena Foster, Kyung Jin Son, Jungmok You, Gulnaz Stybayeva, Dipali Patel, and Alexander Revzin. Cell biology is different in small volumes: Endogenous signals shape phenotype of primary hepatocytes cultured in microfluidic channels. Scientific Reports, 6:srep33980, September 2016. ISSN 2045-2322. doi: 10.1038/srep33980.

5. Joshua Guild, Amranul Haque, Pantea Gheibi, Yandong Gao, Kyung Jin Son, Elena Foster, Sophie Dumont, and Alexander Revzin. Embryonic Stem Cells Cultured in Microfluidic Chambers Take Control of Their Fate by Producing Endogenous Signals Including LIF. STEM CELLS, 34(6):1501–1512, 2016. ISSN 1549-4918. doi: 10.1002/stem.2324.

6. Stefano Giulitti, Enrico Magrofuoco, Lia Prevedello, and Nicola Elvassore. Optimal periodic perfusion strategy for robust long-term microfluidic cell culture. Lab on a Chip, 13(22): 4430–4441, October 2013. ISSN 1473-0189. doi: 10.1039/C3LC50643F.

7. Giovanni G. Giobbe, Federica Michielin, Camilla Luni, Stefano Giulitti, Sebastian Martewicz, Sirio Dupont, Annarosa Floreani, and Nicola Elvassore. Functional differentiation of human pluripotent stem cells on a chip. Nature Methods, 12(7):637–640, July 2015. ISSN 1548-7105. doi: 10.1038/nmeth.3411.

8. Laralynne M. Przybyla and Joel Voldman. Attenuation of extrinsic signaling reveals the importance of matrix remodeling on maintenance of embryonic stem cell self-renewal. Proceedings of the National Academy of Sciences, 109(3):835–840, January 2012. ISSN 0027-8424, 1091-6490. doi: 10.1073/pnas.1103100109.

9. Nishanth V. Menon, Yon Jin Chuah, Bin Cao, Mayasari Lim, and Yuejun Kang. A microfluidic co-culture system to monitor tumor-stromal interactions on a chip. Biomicrofluidics, 8(6): 064118, November 2014. ISSN 1932-1058. doi: 10.1063/1.4903762.

10. Z. Tatárová, J. P. Abbuehl, S. Maerkl, and J. Huelsken. Microfluidic co-culture platform to quantify chemotaxis of primary stem cells. Lab on a Chip, 16(10):1934–1945, May 2016. ISSN 1473-0189. doi: 10.1039/C6LC00236F.

11. Chunhong Zheng, Liang Zhao, Gui’e Chen, Ying Zhou, Yuhong Pang, and Yanyi Huang. Quantitative Study of the Dynamic Tumor–Endothelial Cell Interactions through an Integrated Microfluidic Coculture System. Analytical Chemistry, 84(4):2088–2093, February 2012. ISSN 0003-2700. doi: 10.1021/ac2032029.

12. João T. S. Fernandes, Oldriska Chutna, Virginia Chu, João P. Conde, and Tiago F. Outeiro. A Novel Microfluidic Cell Co-culture Platform for the Study of the Molecular Mechanisms of Parkinson’s Disease and Other Synucleinopathies. Frontiers in Neuroscience, 10, 2016. ISSN 1662-453X. doi: 10.3389/fnins.2016.00511.

13. Qing Zhou, Dipali Patel, Timothy Kwa, Amranul Haque, Zimple Matharu, Gulnaz Stybayeva, Yandong Gao, Anna Mae Diehl, and Alexander Revzin. Liver injury-on-a-chip: Microfluidic co-cultures with integrated biosensors for monitoring liver cell signaling during injury. Lab Chip, 15(23):4467–4478, 2015. ISSN 1473-0197, 1473-0189. doi: 10.1039/C5LC00874C.

14. Marc A. Unger, Hou-Pu Chou, Todd Thorsen, Axel Scherer, and Stephen R. Quake. Monolithic Microfabricated Valves and Pumps by Multilayer Soft Lithography. Science, 288(5463): 113–116, April 2000. ISSN 0036-8075, 1095-9203. doi: 10.1126/science.288.5463.113.

15. Jessica Melin and Stephen R. Quake. Microfluidic Large-Scale Integration: The Evolution of Design Rules for Biological Automation. Annual Review of Biophysics and Biomolecular Structure, 36(1):213–231, June 2007. ISSN 1056-8700, 1545-4266. doi: 10.1146/annurev.biophys.36.040306.132646.

16. Todd Thorsen, Sebastian J. Maerkl, and Stephen R. Quake. Microfluidic Large-Scale Integration. Science, 298(5593):580–584, October 2002. doi: 10.1126/science.1076996.

17. Craig Watson and Samuel Senyo. All-in-one automated microfluidics control system. HardwareX, 5:e00063, April 2019. ISSN 2468-0672. doi: 10.1016/j.ohx.2019.e00063.

18. G. Bar, L. Delineau, A. Häfele, and M. H. Whangbo. Investigation of the stiffness change in, the indentation force and the hydrophobic recovery of plasma-oxidized polydimethylsiloxane surfaces by tapping mode atomic force microscopy. Polymer, 42(8):3627–3632, April 2001. ISSN 0032-3861. doi: 10.1016/S0032-3861(00)00738-2.

19. Stéphane Béfahy, Pascale Lipnik, Thomas Pardoen, Cristiane Nascimento, Benjamin Patris, Patrick Bertrand, and Sami Yunus. Thickness and Elastic Modulus of Plasma Treated PDMS Silica-like Surface Layer. Langmuir, 26(5):3372–3375, March 2010. ISSN 0743-7463. doi: 10.1021/la903154y.

20. Leticia Liste-Calleja, Martí Lecina, Jonatan Lopez-Repullo, Joan Albiol, Carles Solà, and Jordi Joan Cairó. Lactate and glucose concomitant consumption as a self-regulated pH detoxification mechanism in HEK293 cell cultures. Applied Microbiology and Biotechnology, 99(23):9951–9960, December 2015. ISSN 1432-0614. doi: 10.1007/s00253-015-6855-z.

21. Paqui G. Través, Pedro de Atauri, Silvia Marín, María Pimentel-Santillana, Juan-Carlos Rodríguez-Prados, Igor Marín de Mas, Vitaly A. Selivanov, Paloma Martín-Sanz, Lisardo Boscá, and Marta Cascante. Relevance of the MEK/ERK Signaling Pathway in the Metabolism of Activated Macrophages: A Metabolomic Approach. The Journal of Immunology, 188(3):1402–1410, February 2012. ISSN 0022-1767, 1550-6606. doi: 10.4049/jimmunol.1101781.

22. David M. Mosser and Justin P. Edwards. Exploring the full spectrum of macrophage activation. Nature Reviews Immunology, 8(12):958–969, December 2008. ISSN 1474-1741. doi: 10.1038/nri2448.

23. Peter J. Murray, Judith E. Allen, Subhra K. Biswas, Edward A. Fisher, Derek W. Gilroy, Sergij Goerdt, Siamon Gordon, John A. Hamilton, Lionel B. Ivashkiv, Toby Lawrence, Massimo Locati, Alberto Mantovani, Fernando O. Martinez, Jean-Louis Mege, David M. Mosser, Gioacchino Natoli, Jeroen P. Saeij, Joachim L. Schultze, Kari Ann Shirey, Antonio Sica, Jill Suttles, Irina Udalova, Jo A. van Ginderachter, Stefanie N. Vogel, and Thomas A. Wynn. Macrophage Activation and Polarization: Nomenclature and Experimental Guidelines. Immunity, 41(1):14–20, July 2014. ISSN 1074-7613. doi: 10.1016/j.immuni.2014.06.008.

24. Ting Liu, Lingyun Zhang, Donghyun Joo, and Shao-Cong Sun. NF-κB signaling in inflammation. Signal Transduction and Targeted Therapy, 2(1):1–9, July 2017. ISSN 2059-3635. doi: 10.1038/sigtrans.2017.23.

25. Samuel D. Perli, Cheryl H. Cui, and Timothy K. Lu. Continuous genetic recording with self-targeting CRISPR-Cas in human cells. Science, 353(6304):aag0511, September 2016. ISSN 0036-8075, 1095-9203. doi: 10.1126/science.aag0511.

26. Kevin A. Heyries and Carl L. Hansen. Parylene C coating for high-performance replica molding. Lab on a Chip, 11(23):4122, 2011. ISSN 1473-0197, 1473-0189. doi: 10.1039/c1lc20623k.

27. Vijaya Sunkara, Dong-Kyu Park, Hyundoo Hwang, Rattikan Chantiwas, Steven A. Soper, and Yoon-Kyoung Cho. Simple room temperature bonding of thermoplastics and poly(dimethylsiloxane). Lab on a Chip, 11(5):962–965, March 2011. ISSN 1473-0189. doi: 10.1039/C0LC00272K.

28. Martin Zimmermann, Emmanuel Delamarche, Marc Wolf, and Patrick Hunziker. Modeling and Optimization of High-Sensitivity, Low-Volume Microfluidic-Based Surface Immunoassays. Biomedical Microdevices, 7(2):99–110, June 2005. ISSN 1387-2176, 1572-8781. doi: 10.1007/s10544-005-1587-y.

29. Qingming Hu, Yukun Ren, Weiyu Liu, Ye Tao, and Hongyuan Jiang. Simulation Analysis of Improving Microfluidic Heterogeneous Immunoassay Using Induced Charge Electroosmosis on a Floating Gate. Micromachines, 8(7):212, July 2017. doi: 10.3390/mi8070212.

30. Murat Kuscu and Ozgur B. Akan. Modeling convection-diffusion-reaction systems for microfluidic molecular communications with surface-based receivers in Internet of Bio-Nano Things. PLOS ONE, 13(2):e0192202, February 2018. ISSN 1932-6203. doi: 10.1371/journal.pone.0192202.

31. Hasnia Hajji, Lioua Kolsi, Walid Hassen, Abdullah A. A. A. Al-Rashed, Mohamed Naceur Borjini, and Mohamed Ahmed Aichouni. Finite element simulation of antigen-antibody transport and adsorption in a microfluidic chip. Physica E: Low-dimensional Systems and Nanostructures, 104:177–186, October 2018. ISSN 1386-9477. doi: 10.1016/j.physe.2018.07.034.

32. Guoqing Hu, Yali Gao, and Dongqing Li. Modeling micropatterned antigen–antibody binding kinetics in a microfluidic chip. Biosensors and Bioelectronics, 22(7):1403–1409, February 2007. ISSN 0956-5663. doi: 10.1016/j.bios.2006.06.017.

33. Feng-Jen Hsieh, Yen-Wei Chen, Yuen Yung Hui, Chun-Hung Lin, and Huan-Cheng Chang. Quantification and Imaging of Antigens on Cell Surface with Lipid-Encapsulated Fluorescent Nanodiamonds. Micromachines, 10(5), May 2019. ISSN 2072-666X. doi: 10.3390/mi10050304.

34. Kenneth A. Davis, Barnaby Abrams, Sujata B. Iyer, Robert A. Hoffman, and James E. Bishop. Determination of CD4 antigen density on cells: Role of antibody valency, avidity, clones, and conjugation. Cytometry, 33(2):197–205, 1998. ISSN 1097-0320. doi: 10.1002/(SICI)1097-0320(19981001)33:2<197::AID-CYTO14>3.0.CO;2-P.

35. D. Barnett, I. Storie, G. A. Wilson, V. Granger, and J. T. Reilly. Determination of leucocyte antibody binding capacity (ABC): The need for standardization. Clinical & Laboratory Haematology, 20(3):155–164, 1998. ISSN 1365-2257. doi: 10.1046/j.1365-2257.1998.00116.x.

36. J. P. Landry, Yaohuang Ke, Guo-Liang Yu, and X. D. Zhu. Measuring Affinity Constants of 1,450 Monoclonal Antibodies to Peptide Targets with a Microarray-based Label-Free Assay Platform. Journal of immunological methods, 417:86–96, February 2015. ISSN 0022-1759. doi: 10.1016/j.jim.2014.12.011.

37. Ryan A. Kellogg, Rafael Gómez-Sjöberg, Anne A. Leyrat, and Savaş Tay. High-throughput microfluidic single-cell analysis pipeline for studies of signaling dynamics. Nature Protocols, 9(7):1713–1726, July 2014. ISSN 1750-2799. doi: 10.1038/nprot.2014.120.

38. Linas Mazutis, John Gilbert, W. Lloyd Ung, David A. Weitz, Andrew D. Griffiths, and John A. Heyman. Single-cell analysis and sorting using droplet-based microfluidics. Nature Protocols, 8(5):870–891, May 2013. ISSN 1750-2799. doi: 10.1038/nprot.2013.046.

39. Hangrui Liu, Ming Li, Yan Wang, Jim Piper, and Lianmei Jiang. Improving Single-Cell Encapsulation Efficiency and Reliability through Neutral Buoyancy of Suspension. Micromachines, 11(1):94, January 2020. ISSN 2072-666X. doi: 10.3390/mi11010094.

40. Muhsincan Sesen and Graeme Whyte. Image-Based Single Cell Sorting Automation in Droplet Microfluidics. Scientific Reports, 10(1):8736, May 2020. ISSN 2045-2322. doi: 10.1038/s41598-020-65483-2.

41. Diego Correa, Rodrigo A. Somoza, and Arnold I. Caplan. Nondestructive/Noninvasive Imaging Evaluation of Cellular Differentiation Progression During In Vitro Mesenchymal Stem Cell-Derived Chondrogenesis. Tissue Engineering Part A, 24(7-8):662–671, April 2018. ISSN 1937-3341. doi: 10.1089/ten.tea.2017.0125.

42. Paul Lin, Yuan Lin, Donald P. Lennon, Diego Correa, Mark Schluchter, and Arnold I. Caplan. Efficient Lentiviral Transduction of Human Mesenchymal Stem Cells That Preserves Proliferation and Differentiation Capabilities. Stem Cells Translational Medicine, 1(12):886–897, December 2012. ISSN 2157-6564. doi: 10.5966/sctm.2012-0086.

43. David R. Stirling, Madison J. Swain-Bowden, Alice M. Lucas, Anne E. Carpenter, Beth A. Cimini, and Allen Goodman. CellProfiler 4: Improvements in speed, utility and usability. BMC bioinformatics, 22(1):433, September 2021. ISSN 1471-2105. doi: 10.1186/s12859-021-04344-9.

44. Savaş Tay, Jacob J. Hughey, Timothy K. Lee, Tomasz Lipniacki, Stephen R. Quake, and Markus W. Covert. Single-cell NF-κB dynamics reveal digital activation and analogue information processing. Nature, 466(7303):267–271, July 2010. ISSN 1476-4687. doi: 10.1038/nature09145.

45. Ryan A Kellogg, Chengzhe Tian, Tomasz Lipniacki, Stephen R Quake, and Savaş Tay. Digital signaling decouples activation probability and population heterogeneity. eLife, 4: e08931, October 2015. ISSN 2050-084X. doi: 10.7554/eLife.08931.

46. Ryan A. Kellogg and Savaş Tay. Noise Facilitates Transcriptional Control under Dynamic Inputs. Cell, 160(3):381–392, January 2015. ISSN 0092-8674, 1097-4172. doi: 10.1016/j.cell.2015.01.013.

47. Samuel Zambrano, Ilario De Toma, Arianna Piffer, Marco E Bianchi, and Alessandra Agresti. NF-κB oscillations translate into functionally related patterns of gene expression. eLife, 5: e09100, January 2016. ISSN 2050-084X. doi: 10.7554/eLife.09100.

