## Supplementary material for "Multiplexed microfluidic chip for cell co-culture": readme.txt

Files provided are full, 100 mm wafer designs. Both designs are centered around the origin, and document units are in microns.

The control layer has already been scaled up by 1.015x, to account for PDMS shrinking during curing (assuming a curing temperature of 80°C).

The flow layer design has 'flow' and 'flow_posts' layer. They are kept separate for easier subsequent processing, if necessary for your mask writing software / service. We subtracted the 'flow_posts' layer from 'flow' in KLayout to yield a final file with the correct regions filled in.
