## Supplementary material for "Multiplexed microfluidic chip for cell co-culture": readme.txt

Co-culture chip glucose diffusion/consumption model
-----------------------------------------------------

The model uses MATLAB's PDE toolbox; to run it, ensure that you have it installed, and that your MATLAB version is recent enough to support live scripts.

`glucose_2d_time_varying.mlx` contains a full description of the model with step-by-step explanations of its parts, with easy ways to visualize the effect of different parameters like chamber dimensions on the 3D (XYT) results.

`glucose_2d_time_varying_parametric.mlx` includes the exact same code but refactored to run as a function. This is used to run the model several times with different values of a given parameter, and outputs figures C-D of the paper.
