## Supplementary material for "Multiplexed microfluidic chip for cell co-culture": readme.txt

COMSOL immunostaining model
----------------------------

The solutions were cleared to reduce the size of the model. To run the study as-is, right-click Study 1 and select "Compute". To modify any parts of the model, read below for details.

The most important/relevant parts are:

- Global Definitions > Parameters 1. These are all the parameters you may need to vary, including the geometry of the channels, flow velocity, diffusion and binding kinetics, antigen and antibody concentrations etc.
- Component 1 > Definitions > Variables 1. You should *not* need to edit any of this, but they are useful to know of. These variables define the association and dissociation rates.
- Study 1 > Parametric sweep: This is an easy way to check the influence of a parameter on the solution. Select a parameter to vary, specify a list of values, and run the model. You can then export the data (Under Results > Export) and visualize the results in MATLAB with the `process parametric sweep results.mlx` live script.
- Study 1 > Time dependent. The problem is solved in two steps: first, the flow profile is solved using the Stationary solver (since it is time invariant). Then, the diffusion and binding reactions are solved with the time dependent solver. Click on here to change the range of times to look at, for example 0 to 1000 seconds in 1-second steps.
	Note: by default, the solver dynamically adjusts the timescale to speed up computation. This led to some very imprecise results for cases where the reactions were relatively fast. To correct this, I set a maximum time step under `Solver Configurations > Solution 1 > Time-dependent solver 1 > Time Stepping > Maximum step`.

After running the model, you can visualize different results:

- Velocity: A 2D heatmap of velocity within the chamber. Not very exciting but a good sanity check.
- Concentration: A 2D heatmap of bulk antibody concentration. This is best visualized by exporting it (Export > Image sequence export / GIF export) to watch it as an animation. Though you will only see clear differences in concentration if the bulk AB concentration is very low and/or surface antigen concentration is high and/or k_on is very high.
- Surface species concentration: a few line graphs (at different timepoints, as specified under "Data") showing surface concentration vs position on the binding surface. Each line should slope downwards gently (indicating that there are more bound antibodies close to the inlet, where fresh antibodies are coming in), and as time progresses, each line should be higher than the last.
- Average surface concentration vs time: This is the average of the previous data. The surface concentration is averaged over the length of the binding surface to give one concentration value per time point (This is done in `Mean surface antibody concentration`). This is plotted along with the density of available sites. You should see the blue line rise rapidly at first, then asymptotically approach the red line.
- A few of the plots are then automatically generated for the solutions of the parametric sweep such as Velocity 2, Pressure, Concentration,... you can ignore these, or you can use them to visualize the results for a parameter value that you select.
- Export: self-explanatory. Right click to export all of these. Make sure to set the export path for each of these (click on each of them).
