## Supplementary material for "Multiplexed microfluidic chip for cell co-culture": processNfkbData.m

```
function summaryTable = processNfkbData(dataPath, experimentTitle, conditionLabels, displayResults)
    % Process and plot CellProfiler data for the NFkB reporter study

    arguments
        % Path to folder containing the CellProfiler output .csv files
        dataPath char
        % The title to use for graphs
        experimentTitle char = ''
        % Labels to give each condition (optional, will get them from the
        % CellProfiler files if not specified)
        conditionLabels cell = {};
        % If true, the plots of mean cell intensity, and number of positive
        % cells will be shown.
        displayResults {mustBeNumericOrLogical} = 1
    end
   
    nucleusTable = importfile(fullfile(dataPath,"/Nuclei.csv"));
    imageTable = importfile(fullfile(dataPath,"Image.csv"));
    
    imageNumbers = unique(nucleusTable.('ImageNumber'));
    conditions = unique(nucleusTable.('Metadata_Condition'));
    
    summaryTable = table('Size', [length(conditions), 8],...
                         'VariableTypes', {'string', 'double', 'double', 'double', 'double', 'double', 'cell', 'cell'},...
                         'VariableNames', {'Condition', 'Mean Intensity', 'SD Intensity', 'Positive cells', 'Total cells', ...
                                           'Percent positive cells', 'Total cells per image', 'Percent positive cells per image'});
    
    % For each condition, we get the intensities of all cells (in
    % images matching that condition), and calculate the mean and
    % SD.
    
    for cond = 1:length(conditions)
        tempTable = nucleusTable(nucleusTable.('Metadata_Condition') == conditions(cond), :);
        intensities = tempTable.('Intensity_MeanIntensity_mKate2_orig');
        intensity_mean = mean(intensities);
        intensity_sd = std(intensities);
    
        % Count the number of cells in each image of this condition that
        % are positive (significantly above the mean background intensity)
        n_positive = 0;
        images = unique(tempTable.('ImageNumber'));
        % Also keep track of cells per image in case there is variability
        % between chambers
        total_cells_per_img = zeros(1, length(images));
        pct_pos_per_img = zeros(1, length(images));
        for i = 1:length(images)
            % Get background value for that image
            bg_mean = table2array(imageTable(imageTable.('ImageNumber') == images(i), 'Intensity_MeanIntensity_mKate2_masked_inverted'));
            bg_sd = table2array(imageTable(imageTable.('ImageNumber') == images(i), 'Intensity_StdIntensity_mKate2_masked_inverted'));
            threshold = bg_mean+2*bg_sd;
            % Count number of cells above threshold
            intensities_img = table2array(tempTable(tempTable.('ImageNumber') == images(i), 'Intensity_MeanIntensity_mKate2_orig'));
            n_positive = n_positive + sum(intensities_img > threshold);
          
            total_cells_per_img(i) = length(intensities_img);
            pct_pos_per_img(i) = 100*sum(intensities_img > threshold)/length(intensities_img);
        end
    
        row = cond;
        summaryTable.('Condition')(row) = conditions(cond);
        summaryTable.('Mean Intensity')(row) = intensity_mean;
        summaryTable.('SD Intensity')(row) = intensity_sd;
        summaryTable.('Positive cells')(row) = n_positive;
        summaryTable.('Total cells')(row) = length(intensities);
        summaryTable.('Percent positive cells')(row) = 100.0*n_positive/length(intensities);
        summaryTable.('Total cells per image')(row) = {total_cells_per_img};
        summaryTable.('Percent positive cells per image')(row) = {pct_pos_per_img};
    end

    means = summaryTable.('Mean Intensity');
    sds = summaryTable.('SD Intensity');
    positive = summaryTable.('Percent positive cells');
  
    % Get labels for conditions
    if isempty(conditionLabels)
        conditionLabels = cellstr(summaryTable.('Condition'));
    end
    labelStrings = conditionLabels;
    labels = categorical(labelStrings);
    labels = reordercats(labels, labelStrings);
    
    if (displayResults)
        % Plot mean cell intensity for each condition
        figure;
        bar(labels, diag(means), 'stacked');
        hold on;
        er = errorbar(labels, means, sds);
        er.LineStyle = 'none';
        er.Color = 'Black';
        hold off;
        ylabel('Mean reporter intensity per cell [A.U.]');
        title({experimentTitle, 'Mean cell intensity'});
    
        % Plot percentage of positive cells for each condition
        figure;
        bar(labels, diag(positive), 'stacked');
        ylabel('Percent positive cells');
        title({experimentTitle, 'Percent positive'});
    
        % More detail: plot percentage of positive cells for each image, grouped by condition
        c = {'#0072BD','#D95319', '#EDB120','#7E2F8E', '#77AC30', '#4DBEEE', '#A2142F', '#0072BD','#D95319', '#EDB120','#7E2F8E', '#77AC30', '#4DBEEE', '#A2142F'};
        figure;
        plotSpread(summaryTable.('Percent positive cells per image'), 'showMM', 2, 'xNames', labelStrings, 'distributionMarkers', 'o', 'distributionColors', c(1:length(labelStrings)));
        ylabel('Percent positive cells');
        title({experimentTitle, 'Percent positive cells per image'});
    end
end

function dataTable = importfile(filename, dataLines)
    % Modified from function generated by MATLAB data import utility
    
    %% Input handling
    
    % If dataLines is not specified, define defaults
    if nargin < 2
        dataLines = [2, Inf];
    end
    
    %% Set up the Import Options and import the data
    opts = detectImportOptions(filename);
    
    % Specify range and delimiter
    opts.DataLines = dataLines;
    
    % Specify file level properties
    opts.ExtraColumnsRule = "ignore";
    opts.EmptyLineRule = "read";
    
    % Specify variable properties
    opts = setvartype(opts,{'Metadata_Condition'}, 'categorical');
    opts = setvaropts(opts, ["Metadata_Condition"], "EmptyFieldRule", "auto");

    % Import the data
    dataTable = readtable(filename, opts);
end
```
