## Supplementary material for "Multiplexed microfluidic chip for cell co-culture": readme.txt

NFkB reporter study analysis
===============================

This folder contains the CellProfiler pipeline (nfkb_reporter.cpproj) and MATLAB analysis file (display_nfkb_results.mlx) used to process the co-culture experiment images.

The CellProfiler results for the data shown in the paper are included (in exp[1-3]), so the MATLAB script can be run to produce the figure and statistical analysis shown in the paper, as well as a breakdown per experiment.

(Images are not included, to save storage space)
