## Supplementary information for "Multiplexed microfluidic chip for cell co-culture"

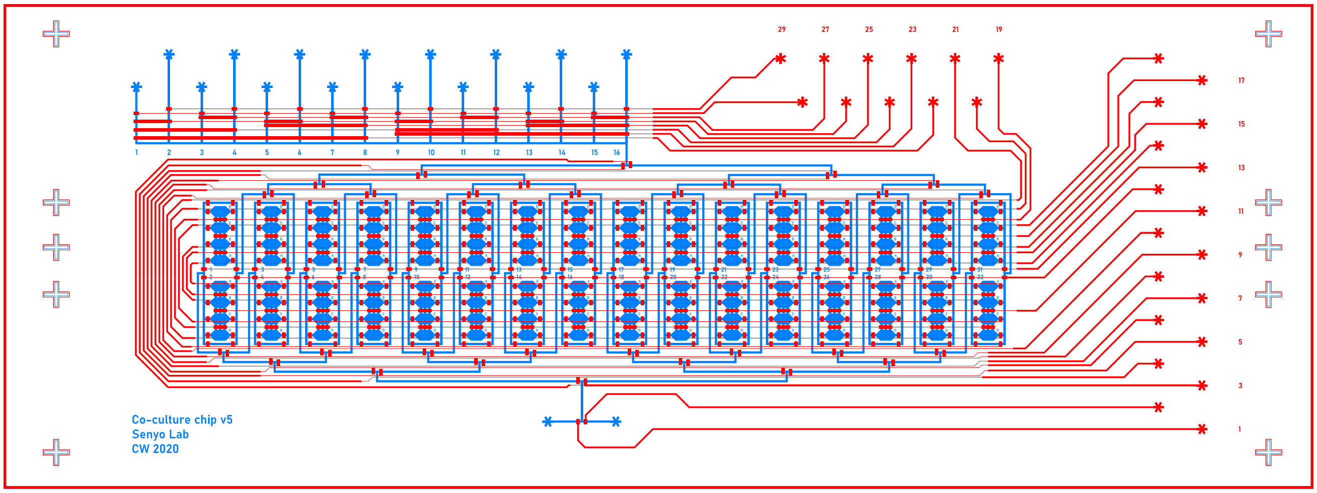


**Fig. S1.** Complete chip design schematic, with flow layer in blue and control layer in red.

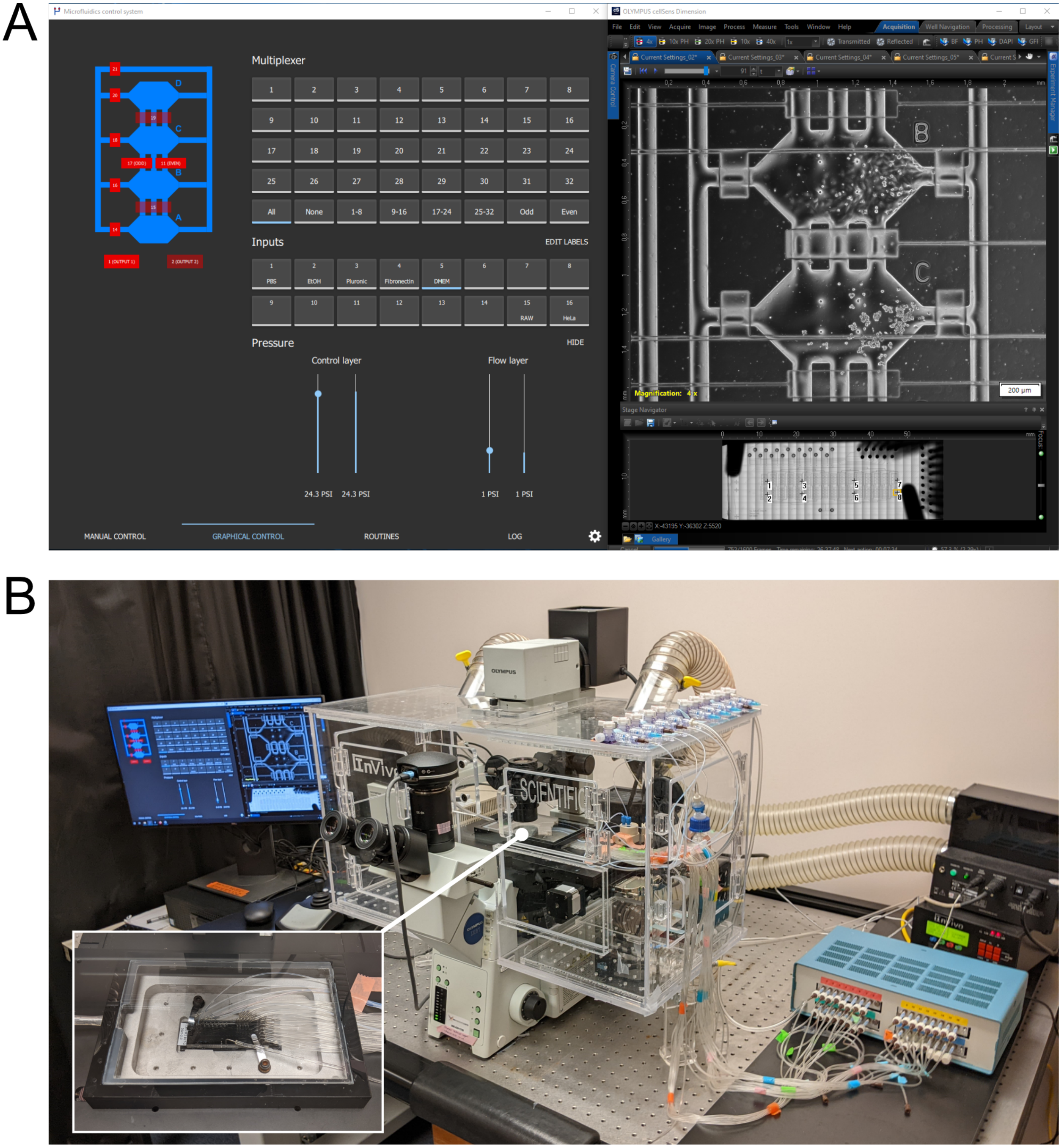


**Fig. S2.** Hardware and software setup **A)** Screenshot of the software in operation. Left: the Qt-based GUI with graphical control showing the live state of the chip and buttons to select multiplexer channel, input, and pressures for the control and flow layers. Some of the most commonly-used multiplexer configurations are provided, while any others are set manually by actuating individual valves (in the manual control screen, not shown). Right: microscope control software (Olympus CellSens) shown here acquiring time-lapse images. **B)** Hardware setup including microscope with heated enclosure and stage-top chamber (inset) for temperature and $CO_2$ control, respectively.

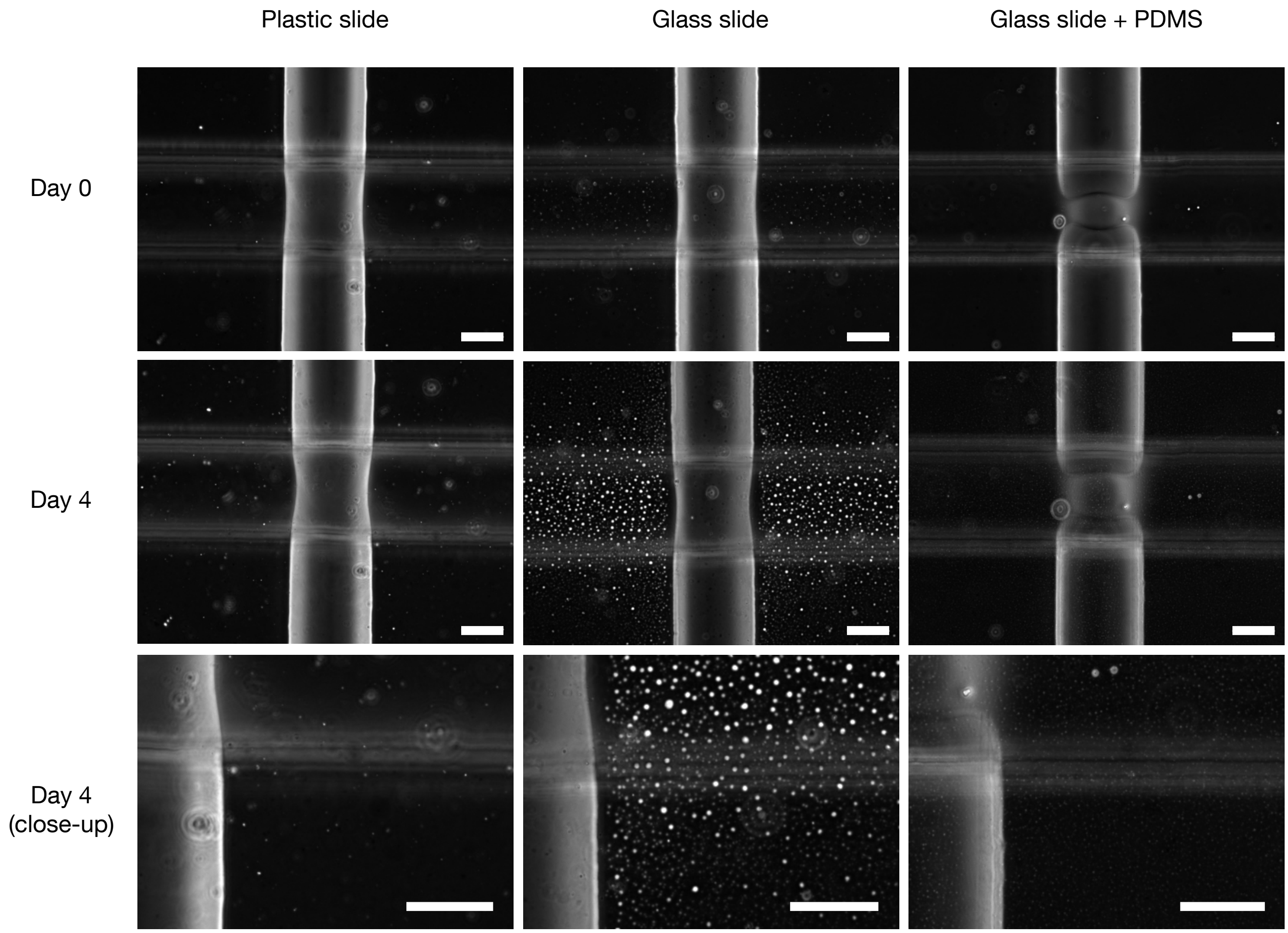


**Fig. S3.** Phase contrast images of a flow channel (vertical) and control channel assembled onto the three different substrates. Water slowly diffuses out of the channels and accumulates on the surface of the glass slides. In the case of PDMS-coated slides, the droplets accumulate below channels and chambers too, introducing artifacts when imaging cells, though the effect is less than with the plasma-treated glass slide. Plastic slides are sufficiently permeable to allow evaporation. In all cases, the water diffusion only affects imaging quality as frequent media replenishment prevents any large changes in osmolarity. Scale bars: 50 µm

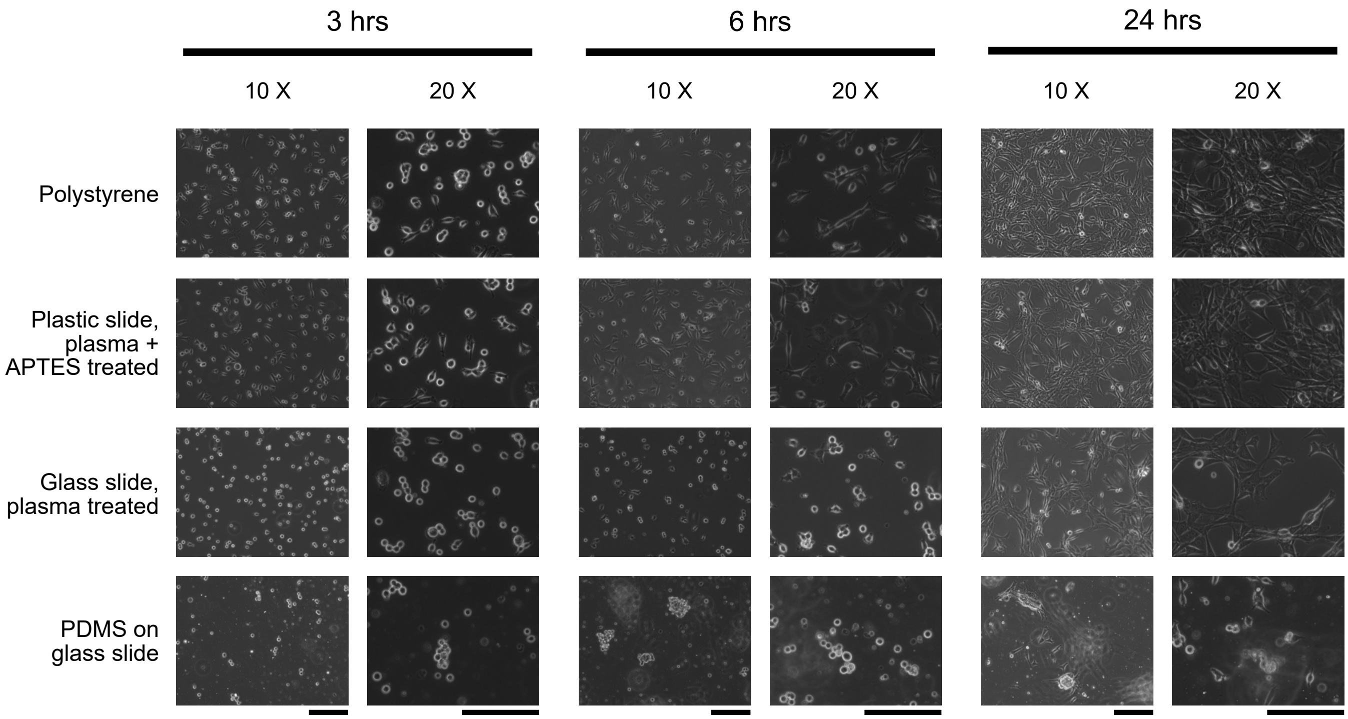


**Fig. S4.** Cell attachment on different substrates. 3T3 fibroblasts were cultured in a 12-well plate, either directly or plated on the different substrates used in the chip. Substrates were prepared in the same way as in chip fabrication. Scale bars: 200 µm.

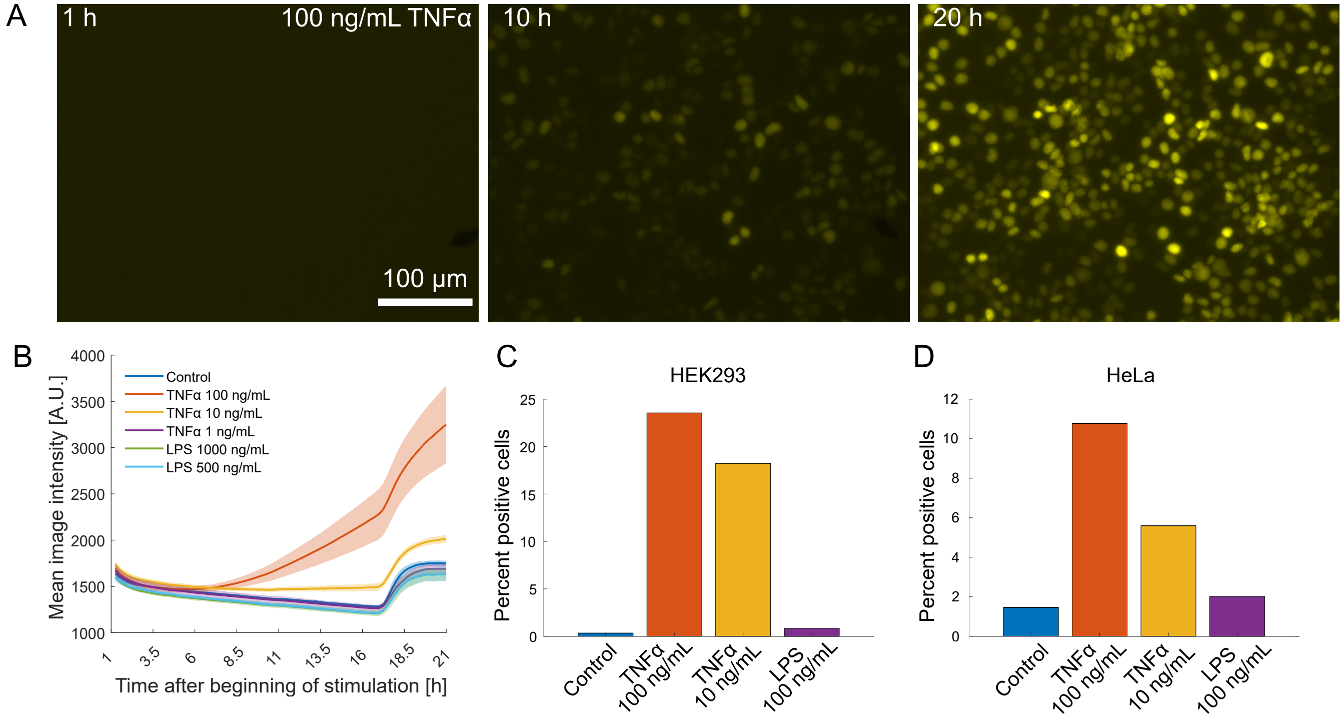


**Fig. S5.** NFκB mKate2 reporter validation. **A)** HEK293Ta treated with 100 ng/mL TNFα, shown at 1, 10 and 20 hours after stimulation onset. **B)** Mean image intensity (raw, unprocessed images) over time of the transduced HEK293Ta. Images were taken every 15 minutes. Shaded areas indicate standard deviation (1 experiment with 3 images per well, 2-4 wells per condition). **C, D)** Endpoint (22 h) analysis of transduced HEK293Ta and HeLa treated with LPS or TNFα (1 experiment with 3 images per well, 2 wells per condition). Cells in C) and D) were imaged only at the 22h timepoint to prevent photobleaching.

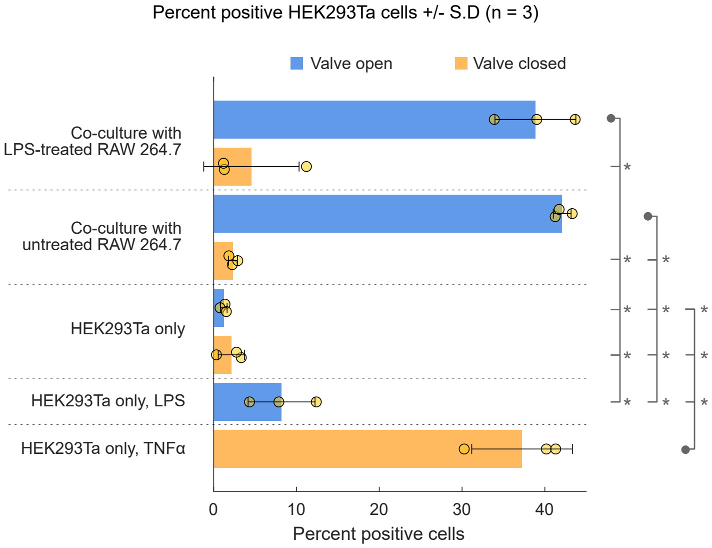


**Fig. S6.** Detailed results of NFκB co-culture assay, showing statistically significant differences between groups ($p < 0.01$). Comparisons are noted with a gray dot marking the group that is significantly different from all groups on the same line as indicated by tick marks and asterisks. Data including analysis script are provided in supplementary file 4.

### Supplementary note 1: Immunocytochemistry protocol

PBS, TX-100 (0.1% in PBS), PFA (paraformaldehyde, 4% in PBS), Stain buffer (0.1% TX-100, 10% goat serum, PBS), primary antibody and secondary antibody solutions are loaded into tygon or PTFE tubing, connected to the chip, and degassed. Then, the reagents are perfused through the culture chamber in the following sequence:

1. PBS, 2 min
2. PFA, 8 min
3. TX-100, 8 min
4. Blocking buffer, 5 min
5. Primary antibody, 15 min
6. Blocking buffer, 2 min
7. Secondary antibody, 10 min
8. (Optional) DAPI stain, 10 min
9. PBS, 2 min

All perfusion steps are done at 2.8 kPa (0.4 psi), through one chamber of all 32 units. Before each step, the reagent is first flushed through the waste channels for 30 seconds at 13.8 kPa (2 psi).
